# TSPDB: A curated resource of tailspike proteins with potential applications in phage research

**DOI:** 10.1101/2024.05.23.595625

**Authors:** Opeyemi U. Lawal, Lawrence Goodridge

**Author notes:** Correspondence: Dr. Opeyemi U. Lawal, Dr. Lawrence Goodridge.

## Abstract

Phages are ubiquitous viruses that drive bacterial evolution through infection and replication within host bacteria. Phage tailspike proteins (TSPs) are key components of phage tail structures, exhibiting polysaccharide depolymerase activity and host specificity. Despite their potential as novel antimicrobials, few TSPs have been fully characterized due to laborious detection techniques. To address this, we present TSPDB, a curated resource for rapid detection of TSPs in genomics and metagenomics sequence data. We mined public databases, obtaining 17,211 TSP sequences, which were filtered to exclude duplicates and partial sequences, resulting in 8,099 unique TSP sequences. TSPDB contains TSPs from over 400 bacterial genera, with significant diversity among them as revealed by the phylogenetic analysis. The top 13 genera represented were Gram-positive, with *Bacillus, Streptococcus*, and *Clostridium* being the most common. Of note, Phage TSPs in Gram-positive bacteria were on average 1 Kbp larger than those in Gram-negative bacteria. TSPDB has been applied in a recent study to screen phage genomes, demonstrating its potential for functional annotation. TSPDB serves as a comprehensive repository and a resource for researchers in phage biology, particularly in phage associated therapy and antimicrobial or biocontrol applications. TSPDB is compatible with bioinformatics tools for *in silico* detection of TSPs in genomics and metagenomic data, and is freely accessible on GitHub and Figshare, providing a valuable resource for the scientific community.

## Background

Bacteriophages (phages) are viruses that infect and replicate within host bacteria and archaea (Chatterjee and Duerkop, 2018; Dion et al., 2020). Phages are the most abundant entities in the biosphere (Dion et al., 2020) and are distributed across different biomes populated by bacterial and archaeal hosts, including the gastrointestinal tract of humans and animals, and oceanic beds (Chevallereau et al., 2022; Clokie et al., 2011). They play a vital role in the rapid evolution and adaptation of their hosts in various environments (Dion et al., 2020).

Phages exhibit high genomic, morphological, and structural diversity, composed of DNA or RNA that can be single-stranded or double-stranded and packaged into a capsid (Dion et al., 2020; Fokine and Rossmann, 2014). The structural form of the capsid was a major feature used in the taxonomic classification of phages until the advent of whole-genome sequencing, which has now become the gold standard for this classification. (Dion et al., 2020; Fokine and Rossmann, 2014; Turner et al., 2023). Phages are broadly classified as tailed or non-tailed, with double-stranded DNA tailed phages constituting about 96% of all known phages (Dion et al., 2020). Phages possess a diverse array of tail structures essential for host recognition, attachment, and penetration, making them important targets in phage therapy research (Fokine and Rossmann, 2014; Gil et al., 2023). Phage infection of its host begins with the recognition of a receptor on the bacterial cell surface for attachment (Dowah and Clokie, 2018; Latka et al., 2017). To penetrate the host cell, phages must overcome various complex barriers on the bacterial cell wall, such as the outer membrane of Gram-negative bacteria and the lipoteichoic acids of Gram-positive bacteria (Chen et al., 2014; Latka et al., 2017). Phages encode virion-associated carbohydrate-degrading enzymes called depolymerases, which are distinct from the endolysins produced by phages during the lysis stage (Knecht et al., 2020; Yan et al., 2014). These depolymerases, encoded by tailspike protein (TSP) genes, recognize, bind, and degrade cell-surface associated polysaccharides, unmasking phage receptors and making them accessible for bacterial infection (Gil et al., 2023; Greenfield et al., 2019; Latka et al., 2017).

Tailspike proteins are integral components of phage tail structures, and their activities as polysaccharide depolymerases are related to host specificity and infectivity (Greenfield et al., 2019). A hallmark of TSPs is their host specificity, high thermostability, resistance to protease treatment, and stability in the presence of high concentrations of urea and sodium dodecyl sulfate (Chen et al., 2014). Phage TSPs possess carbohydrate depolymerase activity and recognize capsule, and lipopolysaccharides (LPS) where they cleave components of the LPS to position the phage towards a secondary membrane receptor during infection (Knecht et al., 2020). TSPs have been observed to decrease bacterial viability, leading to antimicrobial applications. For example, Ayariga and colleagues (Ayariga et al., 2021) demonstrated that the ε34 phage tailspike protein has enzymatic property as a LPS hydrolase and synergizes with Vero Cell culture supernatant in killing *Salmonella* Newington. The ε34 TSP also showed bactericidal efficacy against different *Salmonella* serovars in various matrices (Ibrahim et al., 2023). Miletic and colleagues (Miletic et al., 2016) expressed the receptor binding domain of the Phage P22 Gp9 tailspike protein in plant tissue (*Nicotiana benthamiana*), and demonstrated that, upon oral administration of lyophilized leaves expressing Gp9 TSP to newly hatched chickens, *Salmonella* concentrations were reduced on average by approximately 0.75 log relative to controls. Others have shown that TSPs can be used to control the growth of plant pathogens. For example, expression of the *Erwinia* spp. phage TSP DpoEa1h in transgenic apple and pear plants significantly reduced fire blight (*Erwinia amylovora*) susceptibility, (Malnoy et al., 2005; Roach and Donovan, 2015) likely due to removal of the main virulence factor amylovoran and exposing the *E. amylovora* cells to host plant defenses (Kim et al., 2004). Finally, phage LKA1 TSP exhibits disruptive activity against biofilms while also reducing virulence in *Pseudomonas* in an infection model (Olszak et al., 2017). Collectively, these studies demonstrate the utility of TSPs as novel antimicrobials to control the growth of food and plant-borne pathogens in foods.

Despite the known antimicrobial applications of TSPs, only a few have been fully characterized to date. This could be partly due to the laborious nature of detection techniques, which include plaque assays followed by examination under a transmission electron microscope (TEM) to identify “bulb-like” baseplate structures at the base of phage tails indicative of TSPs (Bhandare et al., 2024; Knecht et al., 2020). The decreasing costs of sequencing and the availability of improved bioinformatics tools have facilitated the construction of large-scale genome and metagenome datasets (Emond-Rheault et al., 2017; Wattam et al., 2014). High-throughput *in silico* detection of TSP-encoding genes in genomic data would not only provide further details regarding the diversity of TSPs in virulent phages but could also be used to identify the presence of TSPs in prophages.

The development of a database for TSPs would further contribute to the understanding of the structure and function of these proteins to harness their potential for diverse applications, such as the development of phage therapy for bacterial infections or phage-based biocontrol of foodborne pathogens, and drug discovery (Brives and Pourraz, 2020; Roach and Donovan, 2015).

Here, we present a high-level curated resource called TSPDB for the rapid detection of tailspike proteins in multiomics sequence data.

## Data and Methodology

### Data Mining and Quality Check

The DDBJ/ENA/GenBank and UniProt databases (Sayers et al., 2022; The UniProt Consortium et al., 2023) were queried for TSPs using search terms commonly associated with tailspike proteins, such as “phage tailspike,” “tail spike proteins,” “phage endopeptidase,” and “phage endorhamnosidase.” Hits were systematically filtered to exclude duplicate results. Nucleotide sequences of TSPs were retrieved from public databases using accession numbers obtained from the database query via NCBI Entrez Programming Utilities (E-utilities) (National Center for Biotechnology Information, 2023)

### Dataset Curation

From this exercise, 17,211 sequences were obtained from the queried public databases. Duplicated sequences were removed using thresholds of ≥ 95% nucleotide similarity and coverage with cd-hit (Li and Godzik, 2006) and Seqkit (Shen et al., 2016), resulting in 9,129 unique TSP sequences (**Figure 1**).

**Figure 1.**
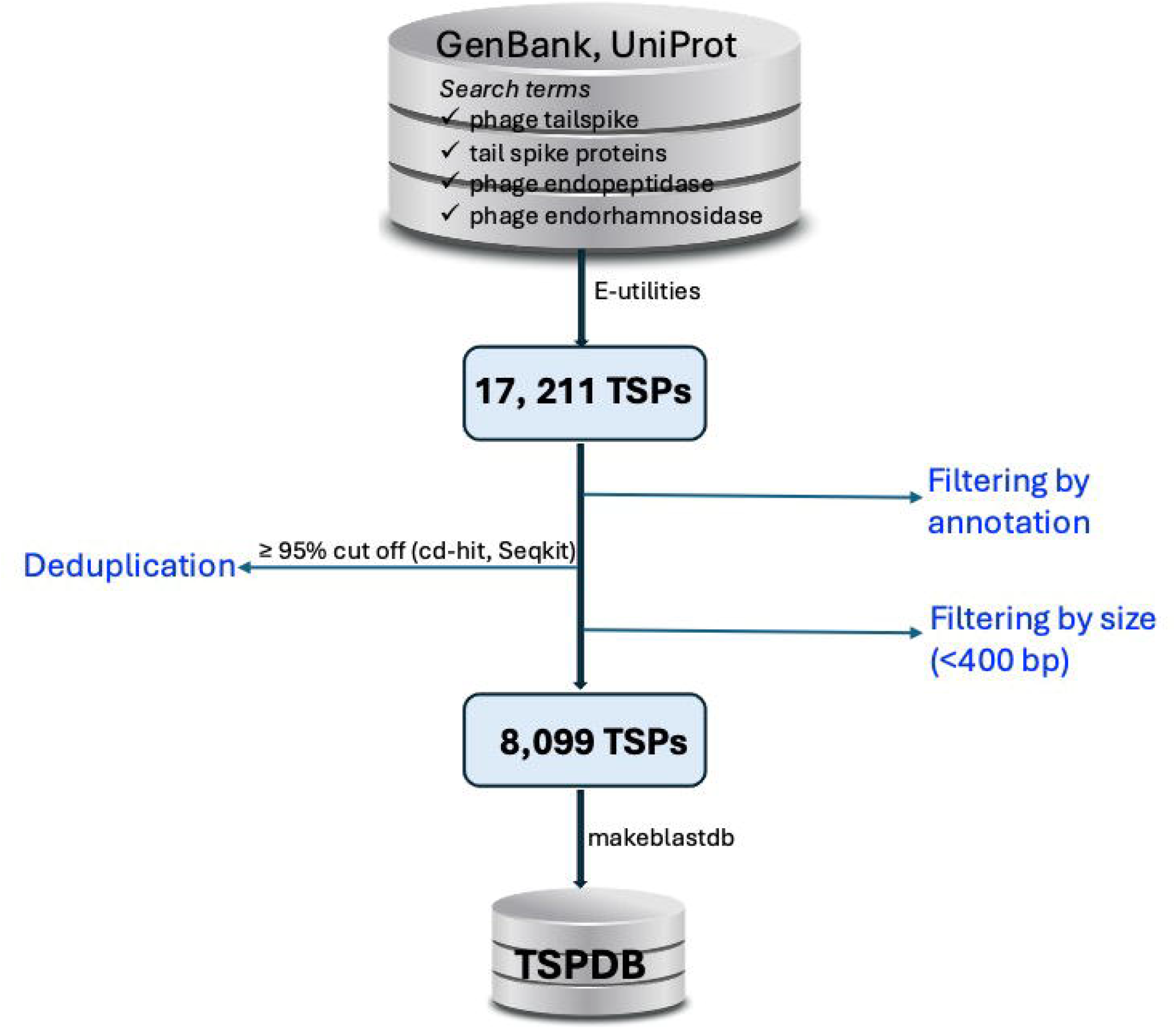
Workflow for the construction of the tailspike protein database (TSPDB).

To assess the sequence length distribution and perform quality checks on unique TSP sequences, Gaussian distribution analysis was conducted. Sequences shorter than 400 bp, which could represent partial or incomplete sequences, were excluded from the dataset. This filtering process resulted in a total of 8,099 unique TSP sequences (**Figure 1**). TSP sequences with a length of ≤10,000 bp were retained to include those originating from Gram-positive bacteria such as *Clostridium* and *Streptococcus*, among others (**Figure 2A**). Further analysis of TSP genes in the TSPDB reveals a significant difference in the sizes of TSPs between Gram-negative and Gram-positive bacteria. Specifically, the average size of TSPs for Gram-negative bacteria is 2,070 bp, while the average size for Gram-positive bacteria is substantially larger, at 3,255 bp (**Figure 2B**).

**Figure 2.**
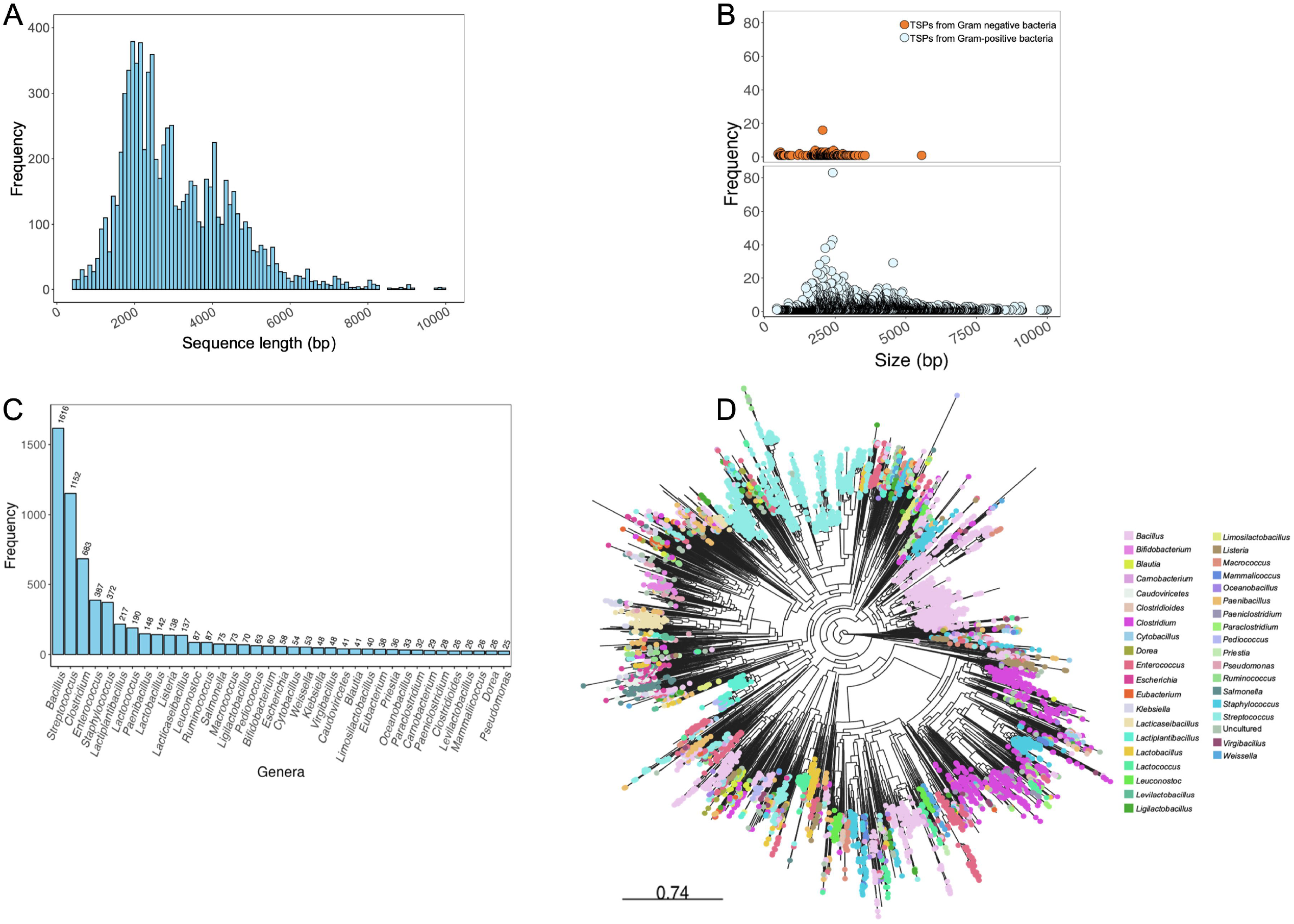
Analysis of Phage tail spike proteins in the TSPDB. (A). Sequence length distribution of genes encoding phage TSPs contained in the TSPDB. (B). Frequency of top 37 genera of host phages carrying TSPs in the TSPDB. (C). Differential TSPs size between Gram-negative and Gram-positive bacteria in the TSPDB. (D). Phylogenetic diversity of the 8,099 TSPs in the TSPDB. Each node represents a unique TSP contained in the TSPDB, with nodes of similar color belonging to the same genera. The top 37 genera are displayed in colour. An interactive version of this figure is accessible through the following link - https://microreact.org/project/7Kv61nb6aRapgGgHpxsNGL-tspdb-v20.

The TSPDB contains TSPs from more than 400 bacterial genera. Among these, the top 13 genera represented were Gram-positive bacteria, with TSPs from *Bacillus* (n=1616) being the most common, followed by *Streptococcus* (n=1152), *Clostridium* (n=683), *Enterococcus* (n=387), and *Staphylococcus* (n=372). Additionally, TSPs from Gram-negative bacterial genera, *Salmonella* (n=75), *Escherichia* (n=58), *Klebsiella* (n=52), and *Pseudomonas* (n=25) were among the top 38 TSPs in the database (**Figure 2C**).

### Diversity of TSPs

To assess the diversity of the 8,099 unduplicated TSP sequences and their suitability for database creation, we employed a phylogeny-based approach. The TSP sequences were aligned using MAFFT v7.453 (Katoh, 2002), and a maximum likelihood tree with 1000 bootstrap replicates for node support was constructed using FastTree v2.1.11 (Price et al., 2010). The resulting phylogenetic tree was visualized using the web-based Microreact visualization tool (Argimón et al., 2016) (**Figure 2D**).

### TSPDB Construction

The deduplicated TSP nucleotide sequences were utilized to construct the TSP database using makeblastdb (Camacho et al., 2009). This database is compatible for use with ABRicate (https://github.com/tseemann/abricate) and other bioinformatics tools equipped with embedded BLAST algorithms, such as BLAST suites and SRST2 (Inouye et al., 2014), among others.

### TSPDB Application

The TSPDB was recently utilized in a study by (Bhandare et al., 2024), where the database was implemented within an ABRicate container to screen for the presence of TSPs in a collection of phage genomes using stringent parameters (≥ 90% identity and ≥ 70% coverage). Overall, the TSPDB contains a vast dataset of diverse TSPs found in phages, making it an essential tool for detecting TSPs within large genomic and metagenomic datasets. Integration of this database into phage detection tools will enhance the functional annotation of these genes. The TSPDB described here will undergo regular updates to include new TSP genes as they become available in public databases.

## Limitations

It is acknowledged that mis-annotation of some TSPs as hypothetical proteins or tail fibers in public databases may have resulted in the omission of certain TSP genes in this study. However, the TSPDB will be continually updated to incorporate additional TSP genes.

## Dataset Description

The TSPDB is freely accessible on GitHub at the following link: https://github.com/yemilawal/Tailspike-proteins or by searching for the title “TSPDB: A curated resource of tailspike proteins with potential applications in phage research” on GitHub. Additionally, accession numbers of genes encoding phage tailspike proteins in TSPDB are available on the GitHub page. A backup version is also available for download on Figshare at https://doi.org/10.6084/m9.figshare.25526323.

## Data Availability Statement

The datasets associated with this study are hosted in online repositories. Details of the repository/repositories and accession numbers can be found in the links provided in the manuscript.

## Author Contributions

OL: Conceptualization, Data curation, Formal analysis, Investigation, Methodology, Validation, Visualization, Writing – original draft and review and editing; LG: Conceptualization, Writing – review and editing, Funding acquisition and Resources

## Publisher’s Note

All claims expressed in this article are solely those of the authors and do not necessarily represent those of their affiliated organizations, or those of the publisher, the editors, and the reviewers. Any product that may be evaluated in this article, or claim that may be made by its manufacturer, is not guaranteed, or endorsed by the publisher.

